# Branched evolution and genomic intratumor heterogeneity in the pathogenesis of cutaneous T-cell lymphoma

**DOI:** 10.1101/804351

**Authors:** Aishwarya Iyer, Dylan Hennessey, Sandra O’Keefe, Jordan Patterson, Weiwei Wang, Gane Ka-Shu Wong, Robert Gniadecki

**Affiliations:** Department of Medicine, University of Alberta, Edmonton, Alberta, Canada; Division of Dermatology, University of Alberta, Edmonton, Alberta, Canada; Department of Oncology, Cross Cancer Institute, University of Alberta, Edmonton, Alberta, Canada; Department of Biological Sciences, University of Alberta, Edmonton, Alberta, Canada; Genesis, Beijing, China; Department of Dermatology, Bispebjerg Hospital, University of Copenhagen, Denmark

## Abstract

Mycosis fungoides (MF) is a slowly progressive cutaneous T-cell lymphoma (CTCL) for which there is no cure. In the early plaque stage the disease is indolent, but development of tumors heralds an increased risk of metastasis and death. Previous research into the genomic landscape of CTCL revealed a complex pattern of >50 driver mutations implicated in more than a dozen of signaling pathways. However, the genomic mechanisms governing disease progression and treatment resistance remain unknown. Building on our previous discovery of the clonotypic heterogeneity of MF, we hypothesized that this lymphoma does not progress in a linear fashion as currently thought, but comprises heterogeneous mutational subclones. We sequenced exomes of 49 cases of MF and identified 28 previously unreported putative driver genes. MF exhibited extensive intratumoral heterogeneity (ITH) of a median of six subclones showing branched pattern of phylogenetic relationships. Stage progression was correlated with an increase in ITH and redistribution of mutations from the stem to the clades. The pattern of clonal driver mutations was highly variable with no consistent mutations between patients. A similar intratumoral heterogeneity was detected in leukemic CTCL (Sézary syndrome). Based on these findings we propose a model of the pathogenesis of MF comprising neutral, divergent evolution of cancer subclones and discuss how ITH impacts the efficacy of targeted drug therapies and immunotherapies of CTCL.

## Introduction

Cutaneous T-cell lymphoma (CTCL) is the most common form of T-cell lymphoma representing 5-10% of the total non-Hodgkin lymphomas.^1,2^ Mycosis fungoides (MF) is the prevalent form of CTCL that initially presents as erythematous, scaly patches and plaques on the skin (stage T1-T2, early lesions) which eventually progresses to advanced lesions, tumors (T3) and erythroderma (T4). Progression to stage T3 is a threshold event during the clinical evolution of MF, associated with a rapid drop in 5-year overall survival from >80% to 44%.^3^ The tumors may appear de novo on the clinically normal skin or may arise within the pre-existing plaque. Therefore, stage T3 patients most often present with a combination of early and late lesions. Thus, MF presents a unique opportunity to study the genetic mechanism of the progression of a T-cell lymphoma and to analyse phyletic relationships between cancer clones in early and advanced stages.

The genomic hallmarks of progressing MF have not been investigated in detail. The majority (84%) of the currently available sequencing data are not derived from MF but from the Sézary syndrome, a leukemic form of CTCL which albeit related, is an entity different from MF.^4–13^ The constellation of mutations in MF is very complex comprising at least 55 potential driver genes, with a considerable variability between patients.^8^ In advanced disease, mutations in the p53 and NFk-B pathways occur frequently and are mutually exclusive, but the survival seems to be similar whichever pathway is affected.^14^ It is also unknown how the pattern of genomic mutations correlate with the phenotype of the lesion.^14^

We hypothesized that lack of a repetitive constellation of mutations in CTCL is a result of the heterogeneity in the mutational processes. It is well documented that in heterogeneous cancers, the bulk sequencing approach will not provide insight into the essential driver genes because most of the detected mutations will be subclonal and only relevant for a subpopulation of malignant cells.^15^ Intratumor heterogeneity (ITH), i.e. existence of genetically different subclones of neoplastic cells within the tumor, has been extensively studied and well documented in different types of solid cancers.^15–17^ ITH turned out to be important in tumor evolution which happens via selection of the fittest subclones. Thus tumors with a high degree of ITH tend to be more aggressive and notoriously difficult to cure because presence of subclones increases the risk of metastasis, facilitates immunological escape and development of resistance to chemotherapy and immunotherapy.^15–17^

The question whether ITH plays a role in the pathogenesis of CTCL has never been addressed before. CTCL is considered to represent a relatively homogenous malignancy in which all neoplastic cells descend from a transformed, mature T-cell in the skin.^18^ We have recently provided evidence supporting an alternative scenario. By analysing the repertoire of T-cell receptor (TCR) sequences in malignant cells in MF, we found that even early lesions, such as patches and plaques, show a considerable level of clonotypic diversity which occurs via seeding of early precursors in the skin.^19–21^ It is conceivable that those malignant clonotypes differ with respect to their mutational history and provide material for ITH in MF.

In this study we were able to confirm the existence of extensive subclonal heterogeneity in MF. We describe differences between early and advanced lesions with respect to the distribution of driver mutations and copy number aberrations and show patterns of branched evolution of cancer genomes during the clinical progression of the disease.

## Material and Methods

### Samples and sequencing

Ethical approval HREBA.CC-16-0820-REN1 was obtained from the Health Research Ethics Board of Alberta, Cancer Committee. Material (4mm punch skin biopsies from lesional skin and 10 ml blood) was collected from 31 consented patients with the diagnosis of mycosis fungoides in stages IA-IVA2 **(Fig 1** and supplementary **Table S1)**. The biopsies and blood were processed for storage as explained in previous methods.^20^ Frozen biopsies were sectioned at 10µm and microdissected to isolate clusters of malignant cells, as previously described in detail.^20^ Peripheral blood mononuclear cells were used as a source of control DNA except for samples MF2 and MF18 for which we did not have matching blood and therefore used microdissected epithelial cells from epidermis as the control. NEBNext® Ultra™ II DNA library prep kit for illumina (cat# E7645S) (New England Biolabs, Massachusetts, United States) was used for DNA library preparation and SSELXT Human All exon V6 +UTR probes (Agilent Technologies, California, United State) were used for the exome and UTR sequence capture. The DNA libraries were sequenced on an Illumina HiSeq 1500 sequencer using paired-end (PE) 150 kit (cat# PE-402-4002) (Hiseq PE rapid cluster kit V2) or NovaSeq 6000 S4 reagent kit 300 cycles (cat# 20012866).

**Figure 1:**
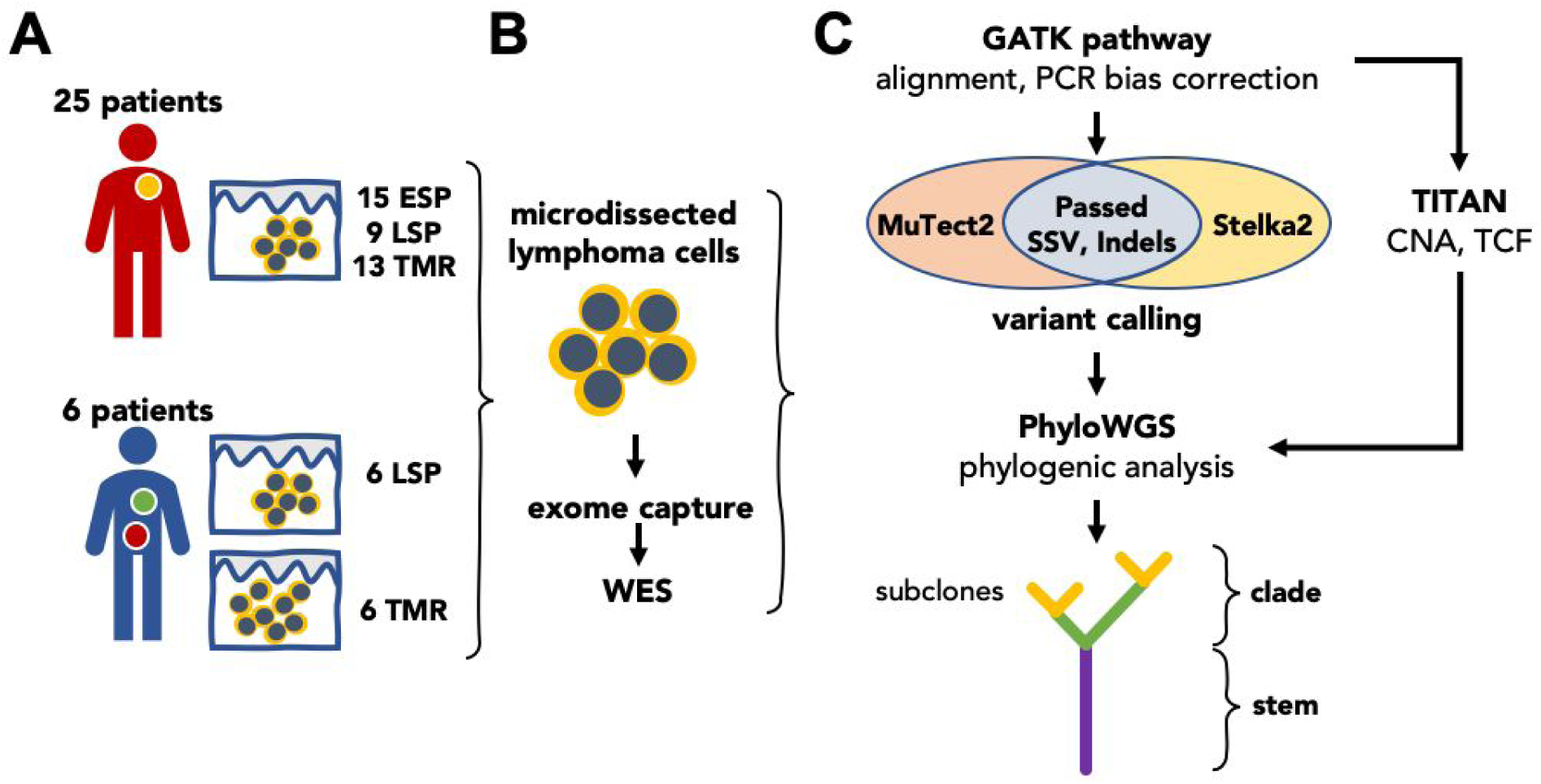
Summary of experimental methods and data analysis. **A.** 49 biopsies was obtained from 31 patients with MF. In 6 patients with tumor stage disease paired biopsies from TMR and LSP were obtained. **B.** Tumor cell clusters microdissected from the lesional skin and matching control tissue (peripheral blood or the epidermis, not shown) were sequenced by WES. **C.** The genetic aberration data (SNV and CNA) was used for the reconstruction of phylogenetic trees of MF. Abbreviations: ESP, early stage plaque, LSP, late stage plaque, TMR, tumor, WES, whole exome sequencing, SV, somatic variants, CNA, copy number aberration, TCF, tumor cell fraction.

### Bioinformatic analysis

The raw fastq files generated from WES were processed through the GATK (version 4.0.10) best practices workflow^22^ and aligned to the hg38 reference genome. Somatic variants (SV), that include Single Somatic Mutations (SSMs) and indels were identified by two different variant callers, MuTect2 (version 2.1)^22,23^ and Strelka2 (version 2.9.10).^24^ The variants filtering as “Passed” by both variant callers were used for downstream analysis. The functional effects of SVs were identified by the Variant Effect Predictor (VEP) (version 95.2).^25^ The Copy Number Aberrations (CNA) and Tumor Cell Fraction (TCF) were identified using TitanCNA (version 1.20.1).^25,26^ PhyloWGS (version 1.0-rc2) was used for phylogenetic analysis to identify the clones and subclones **(Fig 1)**.^25–27^

Sequencing data from previous sequencing studies (supplementary **Table S2)** in CTCL were obtained from public databases and were subjected to the same bioinformatics analysis as described above, with the exception that only MuTect2 was used for variant calling.

### Data availability

The raw sequencing fastq files are submitted to dbGAP with accession number phs001877.v1.p1.

## Results

### Genomic landscape of driver genes in MF

We have decided to revisit the genomics of MF because previous studies have largely focused on Sézary syndrome, a rare leukemia-lymphoma syndrome which although related to MF, is considered to be a separate entity. Only 11% of all previously sequenced CTCL cases were MF and the material for sequencing was the entire skin biopsy which might have introduced an error due to the contribution of mutations from other cells than the lymphoma.^28^ To enrich the material in neoplastic cells, we microdissected clusters of lymphoma cells from 49 skin biopsies from 31 MF patients in various stages (I-IV) (**Supplementary Table S1)** for whole exome sequencing (WES) (**Fig 1**). For the analysis, we decided to group the samples not only by the clinical stage, but also by the morphological features of the biopsied lesion. In stages I-IIA the lesions as per definition were either patches or plaques (abbreviated for the purpose of this paper as ESP, early-stage plaques) but in stages ≥IIB we have distinguished between the biopsies from tumors (TMR) and the plaques (referred to as the late-stage plaques, LSP).

We identified a median of 765 non-synonymous SV in ESP, 1269 in LSP and 2133 in TMR (**Fig 2**), documenting that mutations accumulate during disease progression. These numbers are an order of magnitude higher than was previously found in CTCL (median of 42 non-synonymous mutations^8^). The high number of mutations was likely a result of the high tumor cell fraction (TCF) in our material (**Supplementary Fig S1**) as well as the deeper exome sequencing compared with the previously published data. Among the mutations detected in our material, 265 genes were previously adjudicated as driver genes in cancer^29^ and 56 genes were reported in CTCL.^4–6,8–13^ Current analysis adds 28 additional genes which fulfill the criteria of cancer drivers^29^ that we found to be mutated in >20% of the patients (**Fig 2**, see supplementary **Table S3** for the complete list of the mutated genes). Among those 28 genes, five genes (*ZFHX3, CIC, EP300, PIK3CB* and *HUWE1*) were found in 33-45% of the samples (**Fig 2**). Most mutated genes previously described as drivers, mapped to the already known pathways such as transcription factors (34 genes) followed by chromatin modification (22 genes). The new mutated pathways found here were the Wnt/B Catenin (4 mutated genes), microRNA processing (*DICER1*), protein homeostasis (*HUWE1*) and genome integrity (*POLE, PDS5B*). Lastly, we found that mutations in the tumor suppressor genes dominated over the known oncogenes (supplementary **Fig S2**), analogously to what have previously been reported in other heterogeneous solid tumors.^30,31^

**Figure 2:**
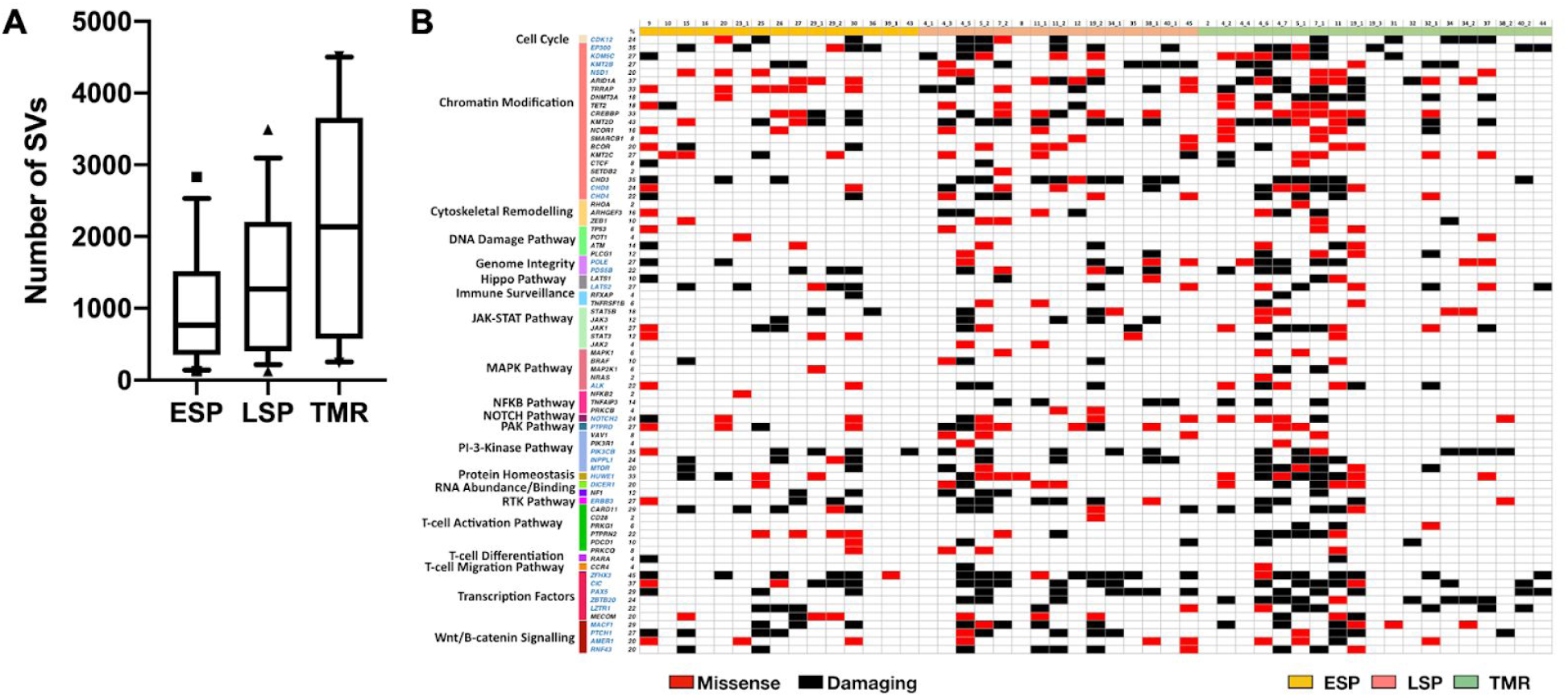
Mutational landscape of putative driver genes in MF. **A.** Number of non-synonymous SVs in samples from ESP, LSP and TMR. Box and whisker plot showing 90^th^ percentile respectively. **B.** Identification of amino acid altering mutations in 75 putative driver genes across 21 different pathways. Black gene symbols annotate the previously reported 47 driver genes in CTCL; the previously unreported 28 potential drivers identified in this study are highlighted in blue. Damaging mutations indicates frameshift mutations, short read insertion and deletion (<6bp), stop gain or stop lost.

We also investigated the CNA profiles for all our samples (**Fig 3A**) and were able to confirm the previously noted amplifications of chromosome 1 and 7 and deletions in chromosome 9 in MF.^32^ The patterns of changes in the CNA profiles were similar for samples TMR and LSP but different from ESP that surprisingly was characterised by larger CNA fragments and increased number of copy number gains across all chromosomes except 6, 9,13, 19 and 21 (**Fig 3A**). We have also analyzed CNA changes in the putative driver genes. We reproduced the previous finding of Choi et al.^13,20^ of the deletion of *TNFAIP3* and found additional genes that were affected in all investigated samples: deletions in *RHOA* (tumor suppressors) and amplifications in the oncogene *BRAF* (**Fig 3B**). In summary, we have found that progression from early stage I to advanced (IIB or higher) was associated with an increased number of non-synonymous SNV, both in the tumors and the plaques. Many of those aberrations affected the potential driver genes and are reported here for the first time.

**Figure 3:**
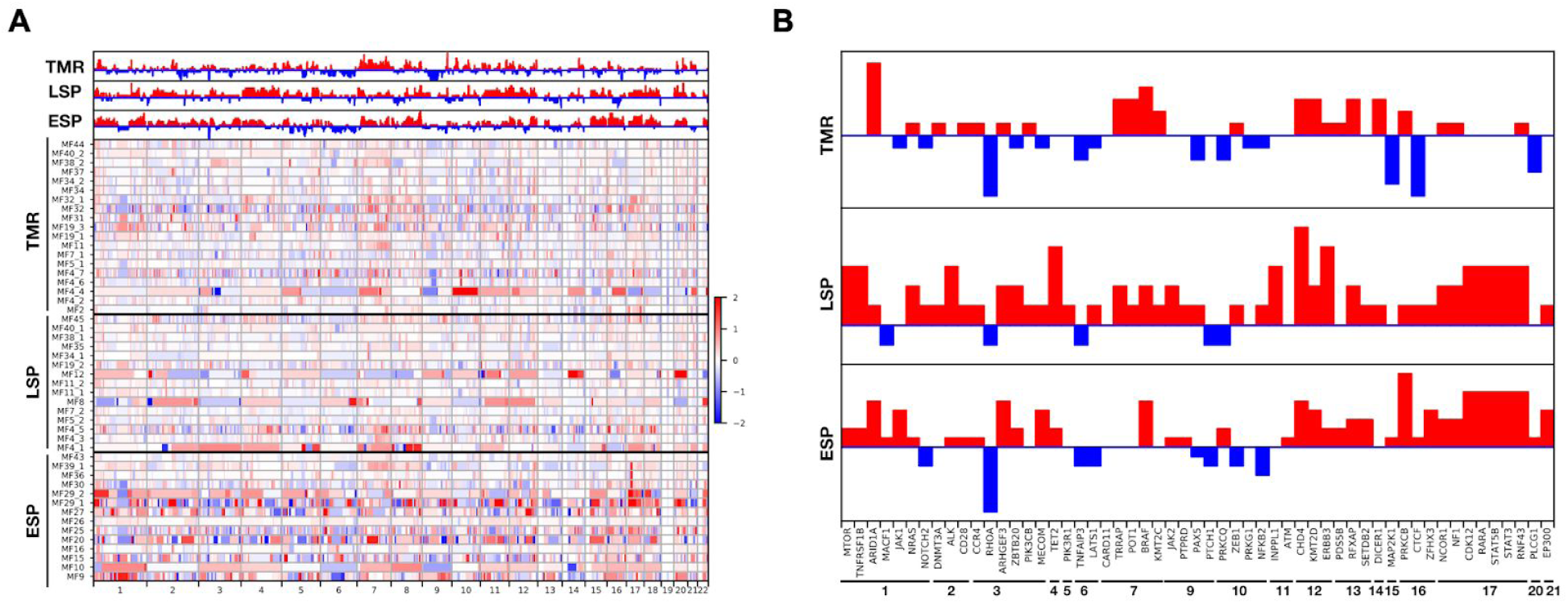
Genomic copy number changes in MF. **A.** Heatmap showing the changes in copy number for the 49 MF samples separated by the type of the lesion (ESP, LSP, TMR). The red bar indicates an amplification and the blue bar indicates a deletion. The number of amplifications or deletions are presented a log2 scale. The bar charts at the top show the difference in number of amplifications (+1 per sample) or deletions (−1 per sample) at each chromosome across lesion types. **B.** Difference in number of amplifications or deletions for putative driver genes in each subgroup of MF samples.

MF is very rich in mutations, both with respect to SNV and CNA. We have therefore asked whether those mutations are clonal or rather a manifestation of the subclonal architecture of MF.

### Intratumor heterogeneity in MF

Genetic aberrations (SVs and CNAs) in solid tumors have often clonal (present in all cells) or subclonal distribution. The subclones may be present in the common stem (“trunk”) of the phylogenetic tree or may occur as a result of branched evolution. In the latter situation the subclonal mutations present in only a subset of cancer cells, often referred to as the clades, or the branches of the tree **(Fig 1)**.^16,33^

To investigate whether MF is characterized by a subclonal structure, we have used a bioinformatic approach where the combined information from SVs and CNA for each of our samples was used to reconstruct a phylogenetic tree. None of the 49 MF samples analyzed here was clonal. We found the median of 6 subclones with a maximum of 9 clones (**Fig 4A** and supplementary **Fig S3**). The TMR samples tended to have more subclones that ESP or LSP and the phylogenetic trees of TMR were also more branched than those of the plaques. The highly branched trees (more than one branching node) were found in 12/19 of TMR (95% CI 40.9%-81.8%) and in 14/27 (52%; 95% CI 33.6-69.8) of the plaques. Only in a minority of cases the phylogenetic tracing showed a linear pattern of subclones (one case of ESP and 2 cases of LSP and TMR) **(Fig 4B**, supplementary **Fig S3)**.

**Figure 4:**
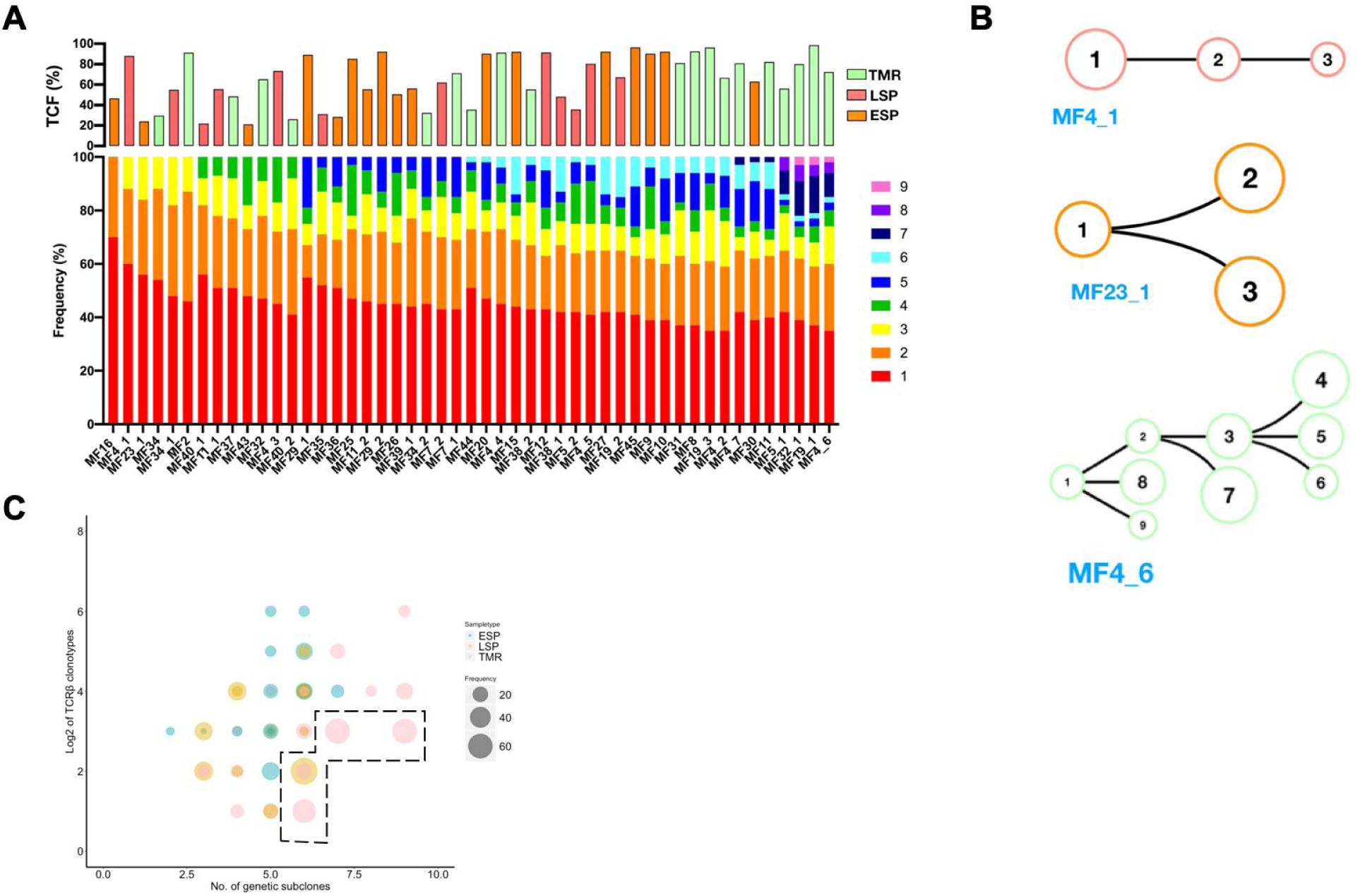
Intratumoral heterogeneity in MF. Combined data from SVs and CNA for each sample was subjected to phylogenetic analysis to identify genetic subclones, as in Fig 1. **A.** Rainbow graph representing the number and frequency of the subclones identified in each sample. The samples are arranged by increasing number of subclones. The top bar graph shows TCF for each sample; the colour of the bars indicate the type of the lesion (ESP, LSP, TMR). **B.** Examples of three major categories of phylogenetic trees: non-branched linear sequence of subclones (upper), simple branched structure with one generation of subclones (middle) and complex structure with several generations of subclones (lower). All phylogenetic trees are shown in supplementary Figure S3. **C.** Bubble plot showing correlation between the number of neoplastic clonotypes and the number of subclones in the samples. The size of the bubble is proportional to the frequency of the first-ranked (the most abundant) clonotype. Dashed line highlights the samples where the first-ranked clonotype had a relative frequency of 60% or higher.

The phylogenic trees reconstructed from the analysis of the distribution of mutations do not necessarily reflect presence of actual cellular clones, defined as a group of identical cells that share a common ancestry. However, being derived from mature T-cells, MF provides an additional opportunity to analyze clonal composition by counting the clonotypes, the unique CDR3 sequences of rearranged *TCRB* genes.^21^ Because *TCRB* locus is rearranged on only one allele and not re-rearranged in mature T-cells, it is possible to calculate the richness and diversity of the repertoire of T-cells. To avoid confusion between different definitions of clones, we will refer to the TCR heterogeneity as the “clonotypic”.

There was a weak, but significant correlation between ITH and clonotypic richness **(Fig 4C)** and between the respective Simpson diversity indices (**supplementary Fig S4**). This suggests that MF is characterized by a divergent evolution, in which the individual T-cell clones accumulate mutations independently of one another. However, we also noticed that the samples in which the most abundant TCRβ clonotype outnumbered the remaining clonotypes (relative frequency ≥60%) also had multiple mutational subclones (>5) (**Fig 4C**). This represented expansion of some neoplastic clones and further branching into multiple mutational subclones (examples of such highly branched phylogenetic trees are MF5_1, MF32_1, MF19_1 and MF4_6 that present with 8-9 subclones, supplementary **Fig S3**).

To further examine whether ITH is present in other types of CTCL, we re-analyzed the sequencing data from 56 samples Sézary syndrome, 13 MF and 8 CTCL not specified, available through data sharing platforms (supplementary **Table S2)**. 7 samples of SS did not present any ITH, whereas the remaining cases demonstrated different degree of ITH ranging from 2 to 9 subclones (median of 4 subclones) (supplementary **Fig S5A**). All MF samples showed ITH similar to that found in our material.

### Subclonal distribution of mutations in MF

We compared the mutational burden in the stem and clades of phylogenetic trees. The distribution of mutational burden changed with disease progression. In most ESPs (11 of 15, 73%), the majority (>50%) of the mutations were concentrated in stem (**Fig 5A**). The situation was reversed in advanced stages. In 73% of LSP and 68% of TMR had >50% mutation in the clades rather than in the stem. The gradual enrichment in the mutations in the clades has been found to be characteristic for divergent evolution^34^, which underscores our conclusion that this is the dominant evolutionary pattern of MF. In only 5 of 49 samples all mutations were concentrated in the stem and these were also the samples with the lowest number of subclones (2-3 subclones) (**Fig 5A**). A slightly higher number of cases with clonal (stem) mutations were found in Sézary syndrome (23%) (supplementary **Fig S5B**), but a direct comparison with our data may be affected by different values of TCF (supplementary **Fig S1**).

**Figure 5:**
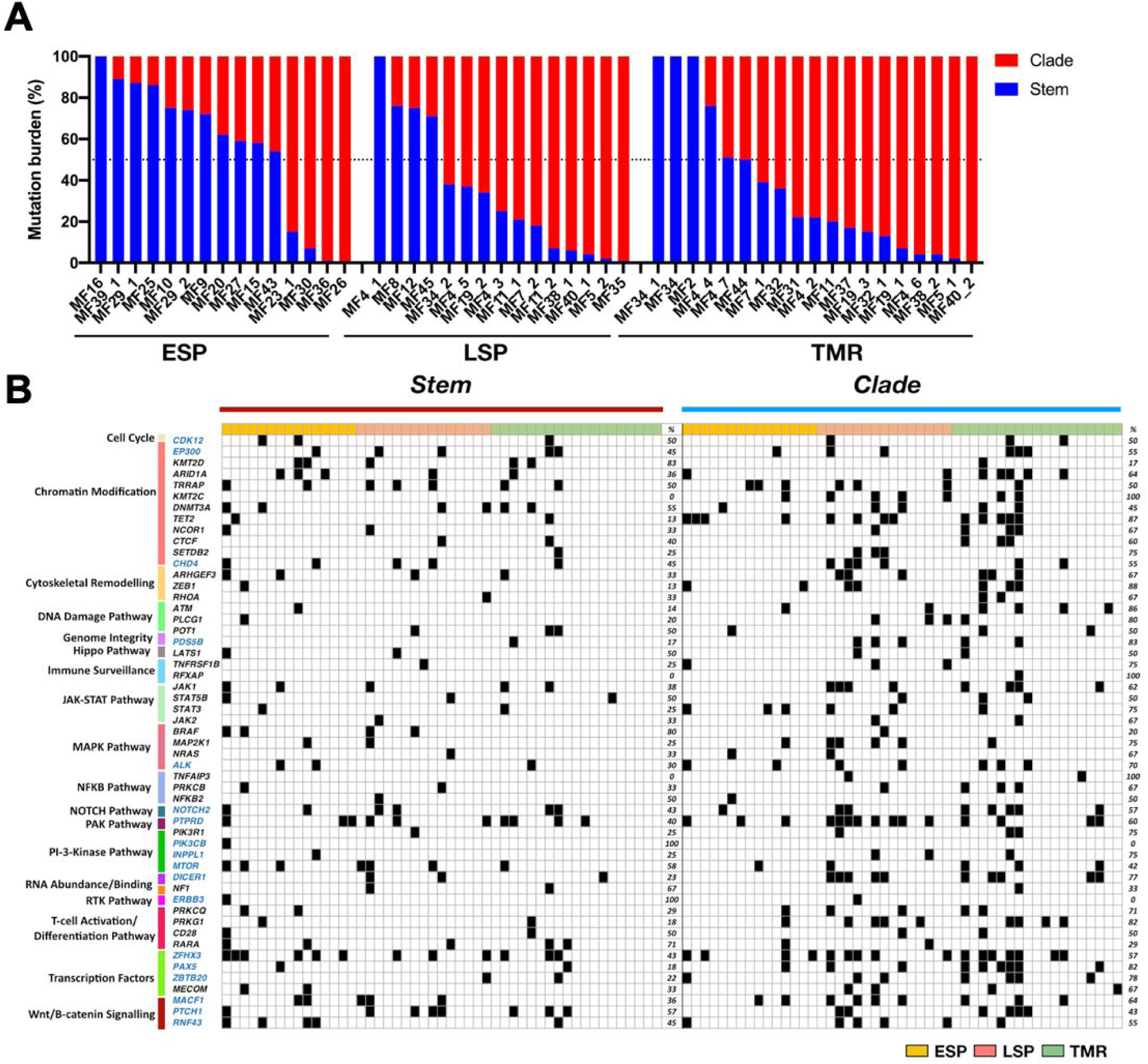
Distribution of mutations in the stem and clades in MF. **A.** Percentage of all SNV mutations in the stem (blue) and clade (red) of the phylogenetic tree. **B.** Mutational landscape of the putative driver genes in the stem and clades of the phylogenetic tree. Black square indicates a function-changing mutation (missense, frameshift, insertions, deletions, stop gain or loss, or variant in 3′ and 5′ UTR). Mutations of the same gene are found both in the stem and the clades signify different position of the mutation.

We have also analyzed the distribution of mutations in the putative driver genes in the stem and clades. Generally, the pattern of driver mutations followed the pattern seen for all SNV with an increasing proportion of mutation in the clades during stage progression (**Fig 5B**). Driver genes such as *CDK12, POT1, LAST1, STAT5B, NFKB2* and *CD28*, representing the pathways of T-cell activation, DNA damage, growth and proliferation, were mutated equally between the stem and clade. However, mutations in some other drivers showed a non-random distribution between the stem and the clade. *PIK3CB* and *ERBB3* were only found in stem whereas *KMT2C, RFXAP* and *TNFAIP3* (essential for chromatin modification and immune surveillance) were only in clades, which as will be discussed below, may have functional importance for the pathogenesis of MF.

### Topologic subclonal heterogeneity in MF

We have previously shown that lesions of MF in the same patient have different clonotypic compositions, a phenomenon which we named “topologic heterogeneity”.^20^ To investigate whether the topologic (interlesional) heterogeneity is also detectable in relation to tumor subclones, we determined the phylogenetic relationships between the subclones in different skin lesions of a single patient. We collected pairs of tumor and plaque biopsies from 6 MF patients and used combined data of SSMs and CNAs from both lesions to map the phylogenetic trees **(Fig 6)**. Two patients (MF5, MF40) had no common ancestral clone shared between the lesions and 3 other patients (MF4, MF7, MF38) had only a single clone shared between the LSP and TMR. Each of the lesion presented an independent phylogenetic branch with multiple subclones (**Fig 6**). We interpret these findings as an additional evidence for divergent evolution of MF witnessing that each lesion evolves in relative isolation from other lesions.

**Figure 6:**
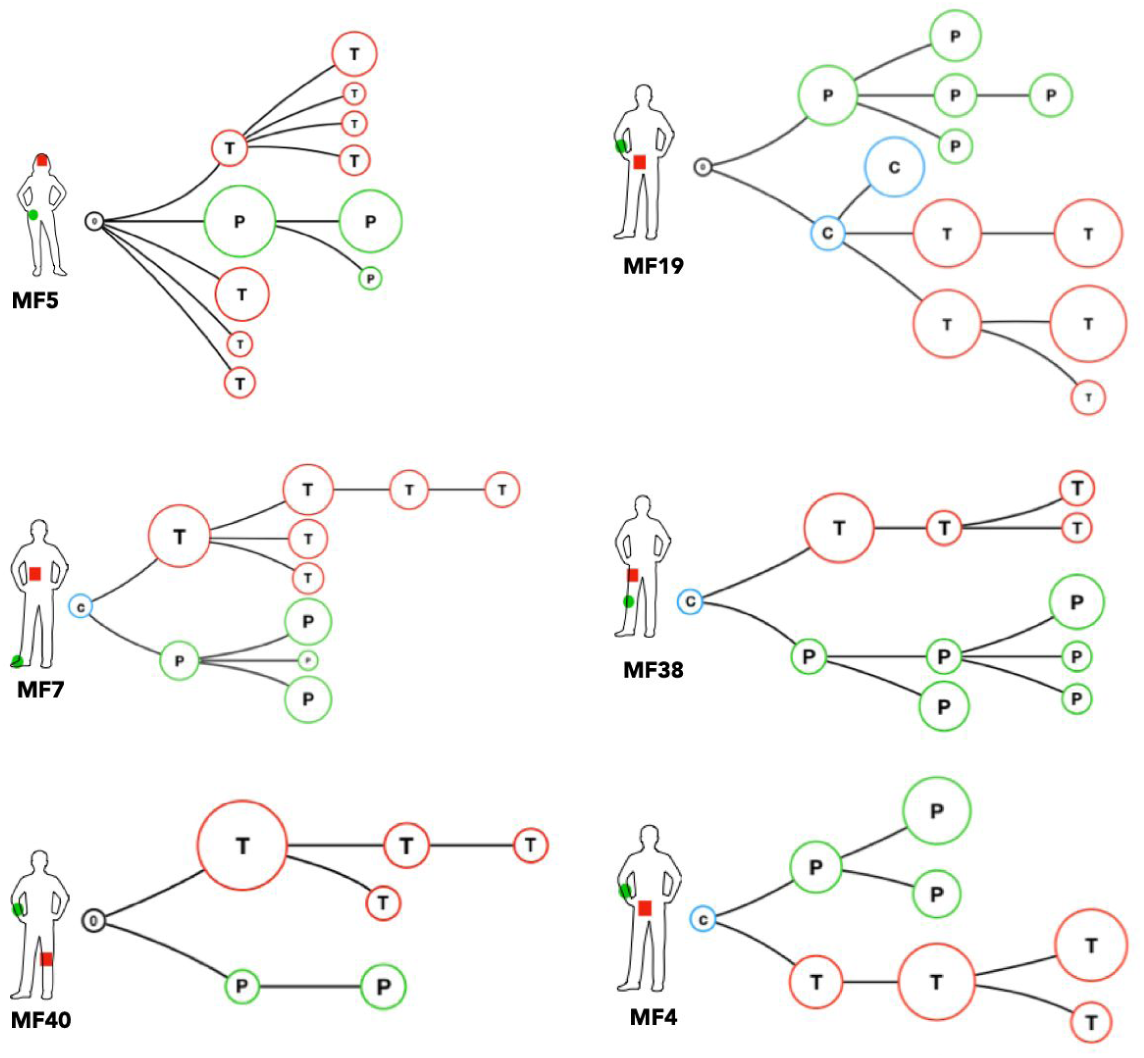
Phylogenetic relationship between different lesions in the same patient: topologic heterogeneity in MF. Pairs of LSP and TMR biopsies were collected from six patients and analyzed as in Fig 4. Red branches represent TMR (T), green branches symbolize the evolution of the LSP (P). The blue circles represent common clones shared by TMR and LSP. The black circle represent a phylogenetic tree without identifiable ancestor clone identifiable.

## Discussion

Intratumor heterogeneity (ITH) refers to the recently described phenomenon of the distribution of somatic mutations in subsets of malignant cells (subclones) rather than being found in all malignant cells (“clonal” mutations). ITH has been documented in solid tumors^16,35^ and in non-Hodgkin lymphomas^36–38^ and is considered to be a genomic manifestation of tumor evolution. ITH arises due to differences in the proliferation and survival between cells bearing different mutations and enhances the evolution by providing material for natural selection where the fittest subclones determine prognosis and resistance to therapy.^15^

Here we demonstrate that MF, previously considered to be a mutationally homogenous lymphoma, is a highly heterogeneous neoplasm composed by multiple subclones. We have found evidence of branched evolution in the majority (92%) of the analyzed MF cases. In addition, subclonal structure was found for the Sézary syndrome, which indicates that ITH is a general feature of CTCL.

Although extensive ITH has been found even in the early stages of MF (T1 in stage IA), disease progression was associated with further accumulation of mutations (SVs and CNA) and increase in ITH. Thus, in contrast to the widely held presumption that progression of MF is caused by a selection and expansion of a single aggressive clone^39^, our data indicate that it is caused by an evolutionary branching leading to enrichment in neoplastic subclones. Several lines of evidence indicate that progression of MF may occur in absence of strong natural selection but happens via divergent evolution which is neutral or only mildly affected by selective pressure on the subclones. The divergent evolution in ITH suggest that each genetic subclone exhibits diversity from the parent clone due to gain or loss of mutations. In addition to the already mentioned hallmarks of neutral tumor evolution such as branched phylogenetic structure and progressive increase in the number of subclones^34^, we have also observed a predicted increase in the number of clade mutations versus stem mutations in advanced stages. Neutral divergent evolution also explains our previous finding why the number of malignant T-cell clonotypes is comparable in early and late stage disease. Malignant T-cell expand and branch into subclones that cohabit the skin niche without evidence of competition and clonal elimination.

We hypothesized that neutral, divergent evolution could explain the resistance of advanced MF to therapy. It has been shown in several types of cancer that ITH is negatively correlated with the sensitivity to chemotherapy or immunotherapy^15,17^ because multiple, genetically diverse malignant subclones provide material for selection of resistant cells. We have explored this question further by analyzing the clonality of putative driver mutations in MF. We added 28 new potential driver mutations to the list of known mutated driver genes in CTCL now including important targetable genes such as *JAK1, JAK3, BRAF, ALK, MTOR*, or *PTCH1*. Unfortunately, we did not find any consistent pattern in the clonal driver mutations which were very heterogeneous and varied vastly from patient to patient. However, we found that certain pathways, in particular chromatin modification and transcription factors were very frequently mutated in the phylogenetic stem in at least one constituent gene in most samples. Based on this observation pathway targeting could be a more promising therapeutic strategy in MF as compared to targeting of specific mutations. Already the drugs that affect chromatin modification mechanisms, such as histone deacetylase blockers, have proved efficacy in MF.^40,41^

Most potential driver mutations in advanced disease are confined to the subclones (clades) which is likely to limit the efficacy of targeted treatment. However, it is worthwhile to mention here that even subclonal mutations may present with drug targeting opportunities if the given subclone is important for the progression of the entire tumor. Cooperativity between distinct subclones was described for some metastasizing tumors^42–44^, and it was proposed that disruption of clonal cooperation might be an interesting therapeutic approach. Especially the targeting of Wnt and Hedgehog seems to be promising^42^, the pathway which we show here to be frequently mutated in MF (genes *MACF1, PTCH1, RNF43*).

Perhaps a better understanding of ITH and the impact of therapy on the subclonal evolution of MF could be gained using multi-omics single cell sequencing of the samples collected before and after the treatment. The subclones detected by bioinformatic reconstruction of phylogenetic trees do not necessarily identify true cellular clones, defined as a group of mutationally identical cells derived from a common ancestor. It has recently been discovered by us and other groups that CTCL is heterogenous at transcriptomics and protein level.^45–48^ Single cell approach would allow for precise mapping of malignant T-cell clones identified by a unique pair of rearranged *TCRA* and *TCRB* sequences to the mutational pattern and gene expression profiles. However, our results identified one potential problem in studies on ITH in multifocal tumor such as MF. By comparing ITH between samples taken from different lesions we found very little phylogenetic relationship between the subclones. In 2/6 cases, the distant lesions did not share any parental clone. Lack of relatedness between discrete lesions could be a result of sampling and missing out the lesion that provides a phylogenetic link. This may indicate that ITH of the totality of MF exceeds significantly the ITH fund in single biopsies. A similar conclusion was reached for ITH of lung cancer by mutational analysis of multiple biopsies.^35^ Driver mutations that were clonal in some biopsies were found to be subclonal in other areas of the tumor. We recognize lack of data from repeated biopsies of multiple skin lesions as a limitation of this study.

Another limitation of our study is the lack of longitudinal follow-up on ITH. We have noted that some phylogenetic trees did not have an identifiable common clone. It is likely that such ancestor clones may become extinct during the progression of the tumor and are no longer detectable, a phenomenon which was described during the evolution of another highly heterogeneous tumor, the glioblastoma.^44^

Finally, we would like to propose a model for the pathogenesis of MF that accounts for the previously found clonotypic heterogeneity^19–21^ and the ITH described here **(Fig 7)**. ITH is readily detectable in ESP, which testifies to the history of mutational tumor evolution before seeding of neoplastic cells in the skin. As the disease progresses, the seeded T-cell clones undergo further mutations and branching into subsequent generations of subclones. Analysis of the branching structure could also confirm our hypothesis that malignant clones from one lesion can re-enter the circulation and seed another lesion. Exchange of malignant T-cell clones between lesions could explain why there is more resemblance between LSP and TMR than between LSP and ESP. A more direct evidence of cancer self-seeding was found in patient MF19 (**Fig 6**) where a subclone shared between the plaque and the tumor was interjected among the branches of the phylogenetic tree.

**Figure 7:**
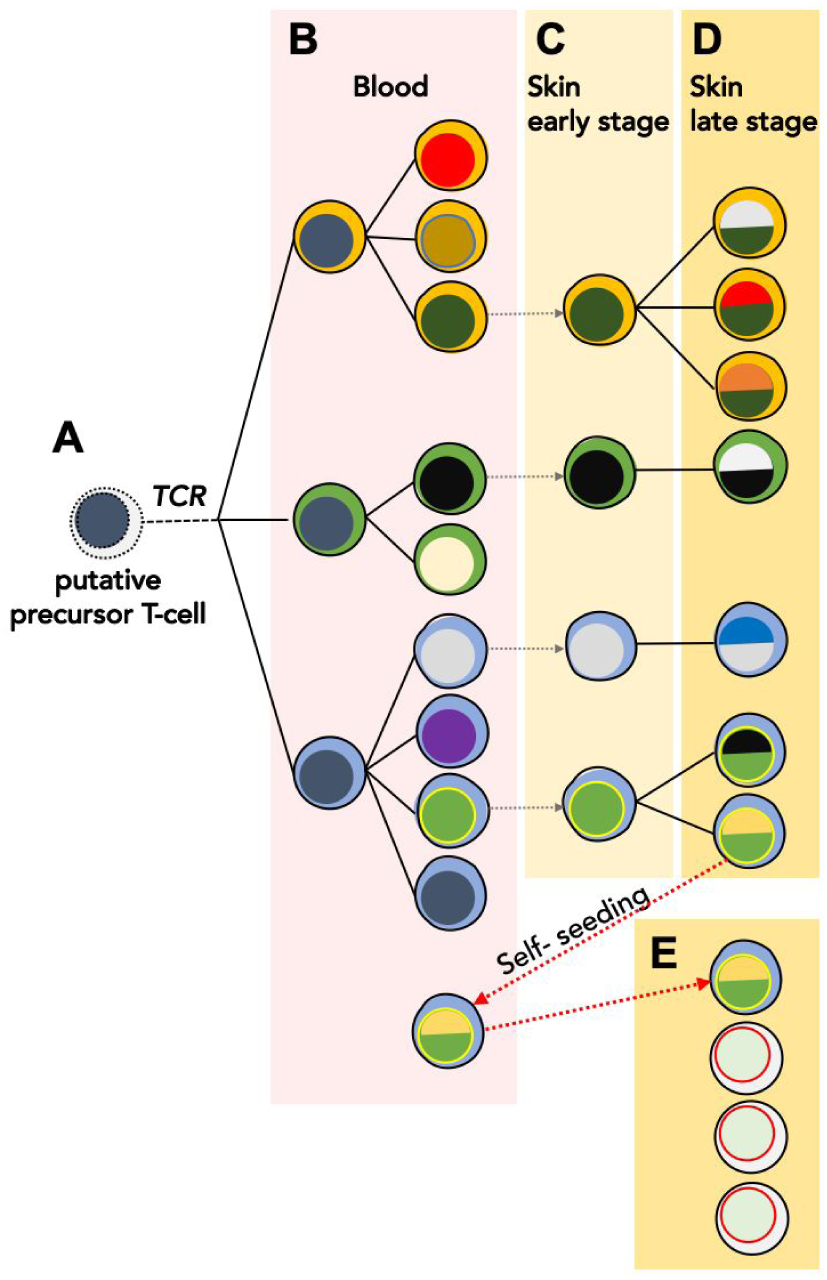
Proposed model of the evolution of MF. The skin lesions of MF are formed by seeding with the circulating malignant T-cell clones which undergo further mutational evolution. It is likely malignant clones originate from an immature T-cell transformed before TCRB rearrangement (**A**) and therefore show clonotypic heterogeneity (highlighted by different colors of the “cytoplasm”). These circulating neoplastic T-cells undergo expansion and accumulate mutations leading to emergence of genetically different malignant subclones (different colors of the “nucleus”) **(B)**. Some of the circulating malignant cells seed into the skin (stippled grey arrows) **(C)** where they proliferate, accumulate further mutations and develop additional subclones as the disease progresses **(D)**. Some subclones may re-enter the circulation and seed other skin lesions (red stippled arrow) further increasing the heterogeneity of the lesions and causing disease progression **(E)**. Solid lines symbolize the phylogenetic relationship between the generations of malignant cells that follow the pattern of divergent, neutral evolution. Based on data in this paper and our previous work.^19–21,49^

## Acknowledgement

We would like to acknowledge Dr. Thomas Salopek, Mrs. Rachel Doucet and the nursing staff of Edmonton Kaye clinic for their help in sample collection. Dr. Hanne Fogh provided excellent care to the patients from the Copenhagen center and helped to collect clinical data. Mrs Vibeke Pless and Mrs Pia Eriksen helped with the collection and shipment of samples. This study was supported by grants from the following sources: Canadian Dermatology Foundation (CDF RES0035718), University Hospital Foundation (University of Alberta), Bispebjerg Hospital (Copenhagen, Denmark), Danish Cancer Society (Kræftens Bekæmpelse R124-A7592 Rp12350) and an unrestricted research grant to R.G. from Department of Medicine, University of Alberta.

## Authors contribution

AI designed the experiments, analyzed the data, wrote the manuscript and submitted data to dbGAP. AI and SO performed the experiments. DH and JP performed the bioinformatic analysis. WW and GW provided input with the technical aspects of the experiments, bioinformatic pipelines, and edited the manuscript. RG designed and supervised the experiments, data analysis and edited the manuscript. All authors approved the final version of this paper.

