## supplementary file.pdf for "Branched evolution and genomic intratumor heterogeneity in the pathogenesis of cutaneous T-cell lymphoma"

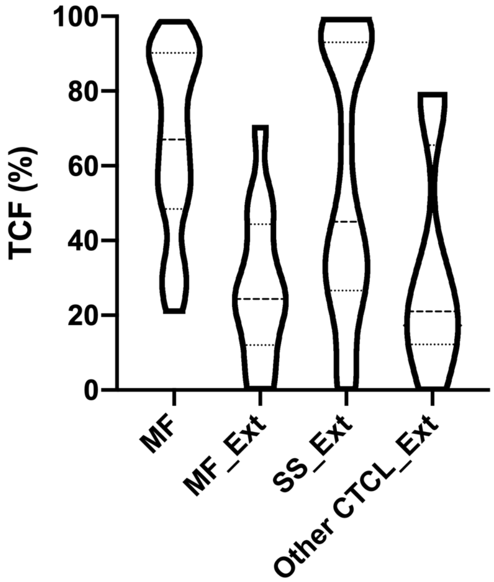


**Supplementary Figure 1: Tumor cell fraction (TCF) in analyzed CTCL samples.** The TCF was calculated for all samples in our study and the samples published in three other CTCL studies (see supplementary Table S2). The samples from the previously CTCL studies were classified as mycosis fungoides (MF), Sézary syndrome (SS) and other CTCL based on the information provided in the original publications (labelled with _Ext for external datasets). The violin plot represents the overall distribution of TCF in the samples.

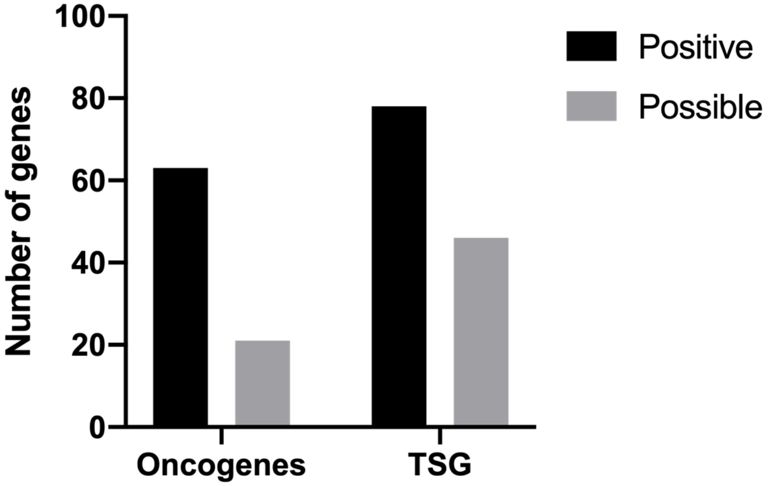


**Supplementary Figure S2: Functional classification of putative driver genes.** Mutations identified in putative driver genes were classified as oncogenes or Tumor suppressor genes (TSG) based on their function with previously reported studies.[30] "Possible" classifies the genes that present a functional ambiguity.

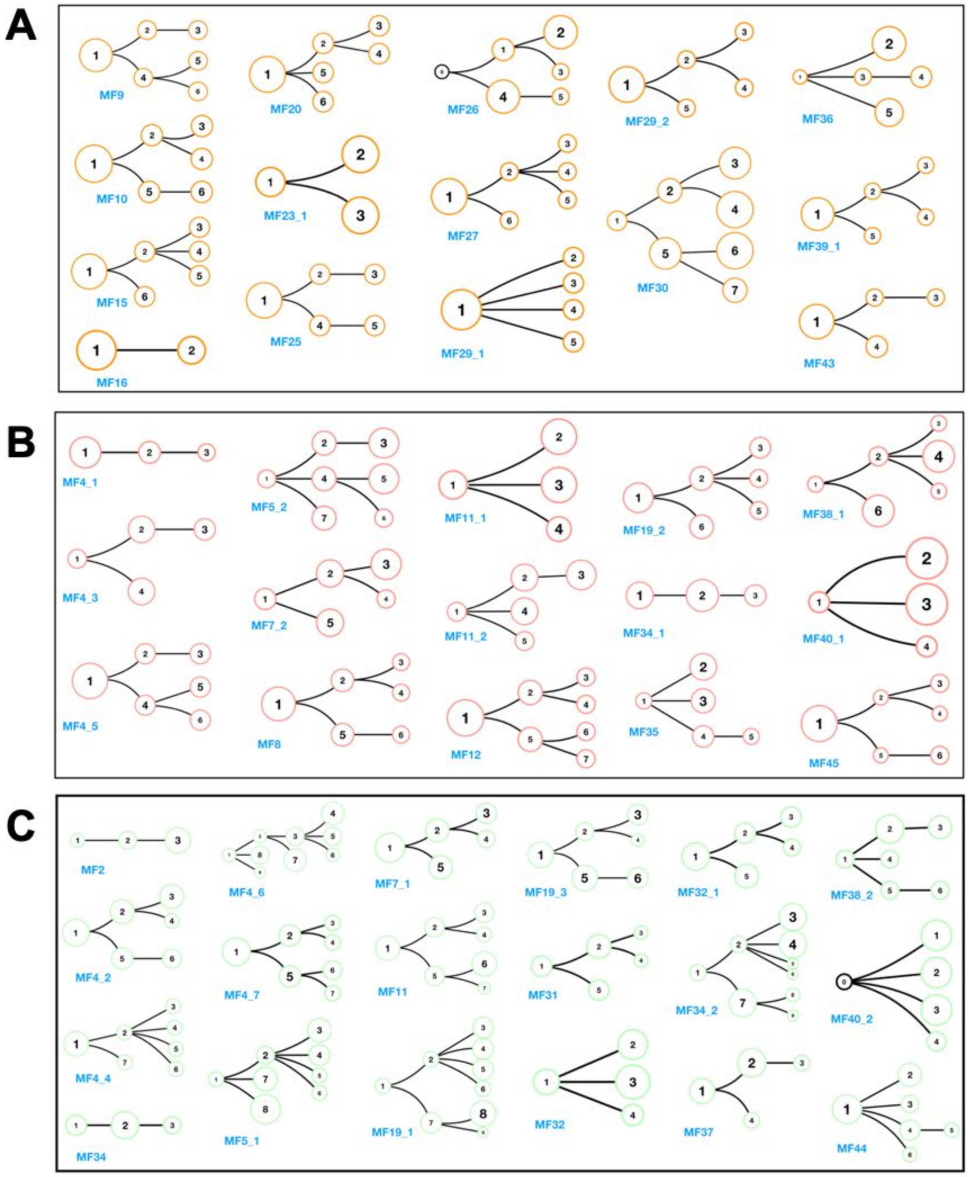


**Supplementary Figure S3: Phylogenetic structures for samples in ESP, LSP and TMR.** (A-C) Presents the phylogenetic trees for every sample in ESP, LSP and TMR respectively. The sample IDs are presented in blue text below each phylogenetic tree.

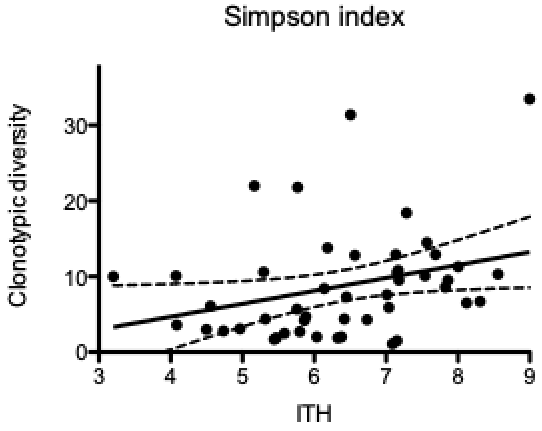


**Supplementary Figure S4: Simpson heterogeneity index for clonotypic and intratumoral heterogeneity (ITH) in MF.** Simpson index for clonotypic heterogeneity was calculated as previously described[1], the ITH Simpson index was calculated and $\frac{1}{D}$ where $D = \frac{N(N-1)}{\sum n(n-1)}$ (*N* - total number of subclones, *n* - the frequency of the individual subclone). A regression line with 95% CI is plotted, $R^2$=0.11, p=0.028.

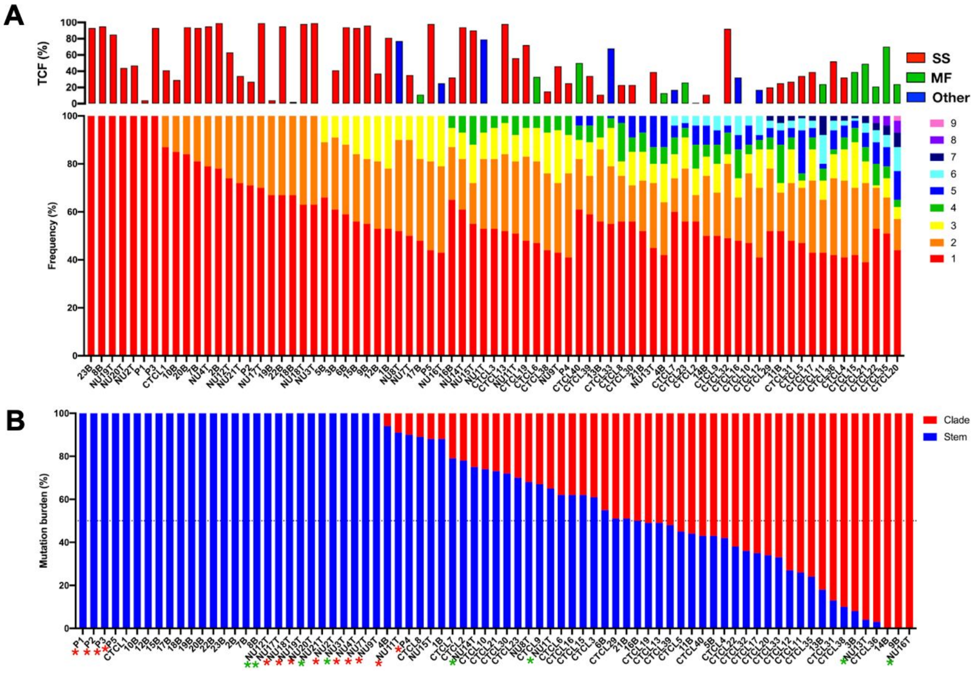


**Supplementary Figure S5: Intratumoral heterogeneity in CTCL.** Data obtained from previous sequencing studies (supplementary Table S2) were analyzed by the same bioinformatic pipeline as in Figure 4A and Figure 5A. **A.** Rainbow graph representing the number and frequency of the subclones identified in each sample. The samples are arranged by increasing number of subclones, followed by the relative frequency of the most abundant subclone. The top bar graph shows TCF for each sample;

the colour of the bars indicate the disease (MF, mycosis fungoides, SS, Sézary syndrome, Other, CTCL unspecified). **B.** Distribution of mutations in the stem and clades.Percentage of all SNV mutations in the stem (blue) and clade (red) of the phylogenetic trees. Asterisk next to the sample names, in red indicate the MF samples and the green indicates CTCL not specified.

**Supplementary Table S1:** Patient characteristics and samples included in the study

| Patient ID ( age [years], sex [M-male, F-female]) | Sample ID | Lesion type | Diagnosis and stage |
|---|---|---|---|
| MF2(83, M) | MF2 | Tumor | Mycosis Fungoides IIB |
| MF4 (69, M) | MF4_1P | Plaque | Mycosis Fungoides IIB |
| | MF4_2T | Tumor | |
| | MF4_3P | Plaque | |
| | MF4_4T | Tumor | |
| | MF4_5P | Plaque | |
| | MF4_6T | Tumor | |
| | MF4_7T | Tumor | |
| MF5 (44, F) | MF5_1T | Tumor | Folliculotropic Mycosis Fungoides IIB |
| | MF5_2P | Plaque | |
| MF7 (62, M) | MF7_1T | Tumor | Mycosis Fungoides IVA2 |
| | MF7_2P | Plaque | |
| MF8 (54, F) | MF8P | Plaque | Folliculotropic Mycosis Fungoides IIIB |
| MF9 (42, F) | MF9P | Plaque | Mycosis Fungoides IA |
| MF10 (56, M) | MF10P | Plaque | Mycosis Fungoides IB |
| MF11 (56, M) | MF11T | Tumor | Mycosis Fungoides IIB |
| | MF11_1P | Plaque | |

| | | | |
|---|---|---|---|
| | MF11_2P | Plaque | |
| MF12 (66, M) | MF12P | Plaque | Mycosis Fungoides IVA2 |
| MF15 (65, M) | MF15P | Plaque | Mycosis Fungoides IB |
| MF16 (68, M) | MF16P | Plaque | Mycosis Fungoides IB |
| MF19 (74, M) | MF19_1T | Tumor | Mycosis Fungoides IIB |
| | MF19_2P | Plaque | |
| | MF19_3T | Tumor | |
| MF20 (70, M) | MF20 | Plaque | Mycosis Fungoides IB |
| MF23_1 (69, F) | MF23_1P | Plaque | Mycosis Fungoides, IA |
| MF25 (48, F) | MF25P | Plaque | Mycosis Fungoides IB |
| MF26 (76, M) | MF26P | Plaque | Mycosis Fungoides IB |
| MF27 (71, M) | MF27P | Plaque | Mycosis Fungoides IA |
| MF29 | MF29_1P | Plaque | Mycosis Fungoides IA |
| | MF29_2P | Plaque | |
| MF30 (62, M) | MF30P | Plaque | Mycosis Fungoides IB |
| MF31 (67, M) | MF31T | Tumor | Folliculotropic Mycosis Fungoides IIB |
| MF32(49, M) | MF32T | Tumor | Mycosis Fungoides IVA |
| | MF32_1T | Tumor | |
| MF34(65, M) | MF34T | Tumor | Mycosis Fungoides IIB |

| | | | |
|---|---|---|---|
| | MF34_1P | Plaque | |
| | MF34_2T | Tumor | |
| MF35 (54, M) | MF35P | Plaque | Folliculotropic Mycosis Fungoides IIB |
| MF36 (64, M) | MF36P | Plaque | Mycosis Fungoides IA |
| MF37 (63, M) | MF37 | Tumor | Mycosis Fungoides IIB |
| MF38(76, M) | MF38_1P | Plaque | Mycosis Fungoides IIB |
| | MF38_2T | Tumor | |
| MF39 (71, M) | MF39_1P | Plaque | Mycosis Fungoides IB |
| MF40(59, F) | MF40_1P | Plaque | Mycosis Fungoides IIB |
| | MF40_2T | Tumor | |
| MF43 (60, M) | MF43P | Plaque | Folliculotropic Mycosis Fungoides IA |
| MF44 (85, M) | MF44T | Tumor | Mycosis Fungoides IIB |
| MF45 (77, M) | MF45P | Plaque | Mycosis Fungoides IIIA |

**Supplementary Table S2:** List of previous CTCL studies used for metaanalysis.

| Study | Sample type | Number of samples |
| --- | --- | --- |
| McGrit.et.al[2] | MF | 5 |
| Choi.et.al[3] | SS | 31 |
| Da Silva Almeida.e[3]t.al[4] | MF, SS and other CTCL | 41 |

**Supplementary Table S3:** Mutations in putative driver presented in <20% of the samples.

| Genes | TMR (%) | LSP (%) | ESP (%) | Gene description | Pathway |
|---|---|---|---|---|---|
| TRAF3 | 2 | 2 | 2 | TNF receptor associated factor 3 | Apoptosis |
| BCL2 | 2 | 0 | 0 | BCL2 apoptosis regulator | Apoptosis |
| BCL2L11 | 2 | 0 | 0 | BCL2 like 11 | Apoptosis |
| CASP8 | 2 | 0 | 0 | caspase 8 | Apoptosis |
| CDK4 | 4 | 0 | 2 | cyclin dependent kinase 4 | Cell cycle |
| BTG2 | 0 | 2 | 0 | BTG anti-proliferation factor 2 | Cell cycle |
| CDKN1A | 2 | 0 | 0 | cyclin dependent kinase inhibitor 1A | Cell cycle |
| ZMYM3 | 6 | 8 | 4 | zinc finger MYM-type containing 3 | Chromatin histone modifiers |
| KANSL1 | 8 | 4 | 4 | KAT8 regulatory NSL complex subunit 1 | Chromatin histone modifiers |
| ARID5B | 6 | 2 | 6 | AT-rich interaction domain 5B | Chromatin histone modifiers |
| KMT2A | 8 | 4 | 2 | lysine methyltransferase 2A | Chromatin histone modifiers |
| SIN3A | 4 | 2 | 2 | SIN3 transcription regulator family member A | Chromatin histone modifiers |
| NIPBL | 6 | 6 | 2 | NIPBL cohesin loading factor | Chromatin other |
| ASXL1 | 8 | 0 | 2 | ASXL transcriptional regulator 1 | Chromatin other |
| AJUBA | 4 | 4 | 0 | ajuba LIM protein | Chromatin other |
| ATF7IP | 2 | 0 | 2 | activating transcription factor 7 interacting protein | Chromatin other |
| H3F3C | 0 | 4 | 0 | H3.5 histone | Chromatin other |
| NPM1 | 0 | 2 | 0 | nucleophosmin 1 | Chromatin other |

| | | | | | |
|---|---|---|---|---|---|
| SMARCA4* | 8 | 4 | 2 | SWI/SNF related, matrix associated, actin dependent regulator of chromatin, subfamily a, member 4 | Chromatin SWI/SNF complex |
| PBRM1 | 8 | 2 | 2 | polybromo 1 | Chromatin SWI/SNF complex |
| ATRX* | 6 | 2 | 2 | ATRX chromatin remodeler | Chromatin SWI/SNF complex |
| SMARCA1 | 2 | 0 | 2 | SWI/SNF related, matrix associated, actin dependent regulator of chromatin, subfamily a, member 1 | Chromatin SWI/SNF complex |
| MSH6 | 2 | 6 | 10 | mutS homolog 6 | Genome integrity |
| POLQ | 6 | 8 | 2 | DNA polymerase theta | Genome integrity |
| SMC1A | 4 | 6 | 2 | structural maintenance of chromosomes 1A | Genome integrity |
| BRCA2 | 2 | 4 | 4 | BRCA2 DNA repair associated | Genome integrity |
| ERCC2 | 0 | 2 | 8 | ERCC excision repair 2, TFIIH core complex helicase subunit | Genome integrity |
| PPM1D | 6 | 4 | 0 | protein phosphatase, Mg2+/Mn2+ dependent 1D | Genome integrity |
| RFC1 | 4 | 4 | 0 | replication factor C subunit 1 | Genome integrity |
| BRCA1 | 2 | 2 | 2 | BRCA1 DNA repair associated | Genome integrity |
| CHEK2 | 2 | 2 | 2 | checkpoint kinase 2 | Genome integrity |
| STAG2 | 4 | 2 | 0 | stromal antigen 2 | Genome integrity |
| ATR | 0 | 2 | 0 | ATR serine/threonine kinase | Genome integrity |
| SETD2* | 6 | 10 | 2 | SET domain containing 2, histone lysine methyltransferase | Histone modification |
| SETBP1 | 6 | 10 | 0 | SET binding protein 1 | Histone modification |
| RNF111 | 8 | 2 | 2 | ring finger protein 111 | Immune signaling |
| IRF6 | 8 | 2 | 0 | interferon regulatory factor 6 | Immune signaling |

| | | | | | |
|---|---|---|---|---|---|
| HLA-A | 0 | 2 | 4 | major histocompatibility complex, class I, A | Immune signaling |
| IL7R | 2 | 2 | 2 | interleukin 7 receptor | Immune signaling |
| HGF | 2 | 2 | 0 | interleukin 6 | Immune signaling |
| B2M* | 0 | 0 | 2 | beta-2-microglobulin | Immune signaling |
| MAP3K4 | 6 | 8 | 0 | mitogen-activated protein kinase kinase kinase 4 | MAPK signaling |
| RPS6KA3 | 2 | 2 | 2 | ribosomal protein S6 kinase A3 | MAPK signaling |
| RRAS2 | 2 | 0 | 0 | RAS related 2 | MAPK signaling |
| IDH2* | 4 | 2 | 0 | isocitrate dehydrogenase (NADP(+)) 2 | Metabolism |
| IDH1 | 4 | 0 | 0 | isocitrate dehydrogenase (NADP(+)) 1 | Metabolism |
| DMD | 8 | 4 | 6 | dystrophin | Other |
| SPTA1 | 10 | 4 | 4 | spectrin alpha, erythrocytic 1 | Other |
| MUC6 | 4 | 6 | 6 | mucin 6, oligomeric mucus/gel-forming | Other |
| GRIN2D | 8 | 2 | 4 | glutamate ionotropic receptor NMDA type subunit 2D | Other |
| SPTAN1 | 8 | 4 | 0 | spectrin alpha, non-erythrocytic 1 | Other |
| FLNA | 2 | 4 | 4 | filamin A | Other |
| MYH9 | 4 | 4 | 2 | myosin heavy chain 9 | Other |
| GABRA6 | 6 | 2 | 0 | gamma-aminobutyric acid type A receptor alpha6 subunit | Other |
| KIF1A | 2 | 2 | 4 | kinesin family member 1A | Other |
| ALB | 2 | 2 | 2 | Fas binding factor 1 | Other |
| TXNIP | 4 | 2 | 0 | thioredoxin interacting protein | Other |
| CNBD1 | 2 | 2 | 0 | cyclic nucleotide binding domain containing 1 | Other |

| | | | | | |
|---|---|---|---|---|---|
| POLRMT | 0 | 2 | 0 | RNA polymerase mitochondrial | Other |
| COL5A1 | 14 | 18 | 14 | collagen type V alpha 1 chain | Other |
| APOB | 12 | 14 | 2 | apolipoprotein B | Other |
| CACNA1A | 10 | 8 | 8 | calcium voltage-gated channel subunit alpha1 A | Other |
| KEL | 10 | 6 | 6 | Kell metallo-endopeptidase (Kell blood group) | Other |
| PTPRC | 8 | 6 | 4 | protein tyrosine phosphatase receptor type C | Other signaling |
| ARHGAP35 | 4 | 8 | 4 | Rho GTPase activating protein 35 | Other signaling |
| GNAS | 10 | 2 | 4 | GNAS complex locus | Other signaling |
| PTPDC1 | 6 | 4 | 2 | protein tyrosine phosphatase domain containing 1 | Other signaling |
| PTPN11 | 10 | 2 | 0 | protein tyrosine phosphatase non-receptor type 11 | Other signaling |
| CDH1 | 6 | 0 | 4 | cadherin 1 | Other signaling |
| KEAP1 | 2 | 4 | 2 | kelch like ECH associated protein 1 | Other signaling |
| SOS1 | 0 | 6 | 2 | SOS Ras/Rac guanine nucleotide exchange factor 1 | Other signaling |
| LEMD2 | 6 | 0 | 0 | LEM domain containing 2 | Other signaling |
| NF2 | 4 | 2 | 0 | neurofibromin 2 | Other signaling |
| PLCB4 | 4 | 0 | 2 | phospholipase C beta 4 | Other signaling |
| GNA11 | 2 | 0 | 2 | G protein subunit alpha 11 | Other signaling |
| GNA13 | 2 | 2 | 0 | G protein subunit alpha 13 | Other signaling |
| GNAQ | 2 | 0 | 2 | G protein subunit alpha q | Other signaling |
| PLXNB2 | 0 | 2 | 2 | plexin B2 | Other signaling |

| | | | | | |
|---|---|---|---|---|---|
| RHOB | 2 | 0 | 2 | ras homolog family member B | Other signaling |
| DIAPH2 | 0 | 2 | 0 | diaphanous related formin 2 | Other signaling |
| GPS2 | 0 | 2 | 0 | G protein pathway suppressor 2 | Other signaling |
| MAP2K4 | 0 | 0 | 2 | mitogen-activated protein kinase kinase 4 | Other signaling |
| PIM1* | 2 | 0 | 0 | Pim-1 proto-oncogene, serine/threonine kinase | Other signaling |
| PRKAR1A | 2 | 0 | 0 | protein kinase cAMP-dependent type I regulatory subunit alpha | Other signaling |
| RAC1 | 2 | 0 | 0 | Rac family small GTPase 1 | Other signaling |
| FAT1 | 22 | 10 | 6 | FAT atypical cadherin 1 | Other signaling |
| PIK3CG* | 8 | 4 | 4 | phosphatidylinositol-4,5-bisphosphate 3-kinase catalytic subunit gamma | PI3K signaling |
| PIK3R2 | 6 | 0 | 0 | phosphoinositide-3-kinase regulatory subunit 2 | PI3K signaling |
| AKT1 | 2 | 0 | 2 | AKT serine/threonine kinase 1 | PI3K signaling |
| PPP2R1A | 0 | 2 | 2 | protein phosphatase 2 scaffold subunit Aalpha | PI3K signaling |
| PIK3CA | 2 | 0 | 0 | phosphatidylinositol-4,5-bisphosphate 3-kinase catalytic subunit alpha | PI3K signaling |
| USP9X | 0 | 6 | 8 | ubiquitin specific peptidase 9 X-linked | Protein homeostasis/ubiquitination |
| CYLD | 6 | 2 | 2 | CYLD lysine 63 deubiquitinase | Protein homeostasis/ubiquitination |
| CUL1 | 6 | 0 | 2 | cullin 1 | Protein homeostasis/ubiquitination |
| EEF2 | 2 | 0 | 4 | eukaryotic translation elongation factor 2 | Protein homeostasis/ubiquitination |

| | | | | | |
|---|---|---|---|---|---|
| BAP1 | 0 | 4 | 0 | BRCA1 associated protein 1 | Protein homeostasis/ubiquitination |
| FBXW7 | 2 | 2 | 0 | F-box and WD repeat domain containing 7 | Protein homeostasis/ubiquitination |
| SPOP | 0 | 4 | 0 | speckle type BTB/POZ protein | Protein homeostasis/ubiquitination |
| VHL | 0 | 4 | 0 | von Hippel-Lindau tumor suppressor | Protein homeostasis/ubiquitination |
| CUL3 | 0 | 2 | 0 | cullin 3 | Protein homeostasis/ubiquitination |
| EEF1A1 | 2 | 0 | 0 | eukaryotic translation elongation factor 1 alpha 1 | Protein homeostasis/ubiquitination |
| ZFP36L2 | 4 | 8 | 4 | ZFP36 ring finger protein like 2 | RNA abundance |
| DHX9 | 2 | 8 | 4 | DExH-box helicase 9 | RNA abundance |
| NUP93 | 8 | 4 | 2 | nucleoporin 93 | RNA abundance |
| CSDE1 | 6 | 2 | 2 | cold shock domain containing E1 | RNA abundance |
| NUP133 | 2 | 6 | 0 | nucleoporin 133 | RNA abundance |
| RBM10 | 2 | 4 | 2 | RNA binding motif protein 10 | RNA abundance |
| ZC3H12A | 2 | 4 | 2 | zinc finger CCCH-type containing 12A | RNA abundance |
| SCAF4 | 2 | 4 | 0 | SR-related CTD associated factor 4 | RNA abundance |
| DDX3X* | 2 | 0 | 0 | DEAD-box helicase 3 X-linked | RNA abundance |
| XPO1 | 0 | 2 | 0 | exportin 1 | RNA abundance |
| ZFP36L1 | 0 | 0 | 2 | ZFP36 ring finger protein like 1 | RNA abundance |

| | | | | | |
|---|---|---|---|---|---|
| ERBB2 | 8 | 2 | 4 | erb-b2 receptor tyrosine kinase 2 | RTK signaling |
| EGFR | 4 | 8 | 0 | epidermal growth factor receptor | RTK signaling |
| EPHA3 | 6 | 0 | 6 | EPH receptor A3 | RTK signaling |
| FGFR2 | 4 | 4 | 4 | fibroblast growth factor receptor 2 | RTK signaling |
| PDGFRA | 6 | 4 | 2 | platelet derived growth factor receptor alpha | RTK signaling |
| EPHA2 | 4 | 2 | 4 | EPH receptor A2 | RTK signaling |
| FGFR1 | 4 | 6 | 0 | fibroblast growth factor receptor 1 | RTK signaling |
| FLT3 | 4 | 4 | 0 | fms related tyrosine kinase 3 | RTK signaling |
| MET | 4 | 2 | 2 | MET proto-oncogene, receptor tyrosine kinase | RTK signaling |
| ERBB4 | 2 | 4 | 0 | erb-b2 receptor tyrosine kinase 4 | RTK signaling |
| KIT | 2 | 4 | 0 | KIT proto-oncogene, receptor tyrosine kinase | RTK signaling |
| RET | 2 | 4 | 0 | ret proto-oncogene | RTK signaling |
| RIT1 | 2 | 0 | 0 | Ras like without CAAX 1 | RTK signaling |
| SRSF2 | 8 | 8 | 0 | serine and arginine rich splicing factor 2 | Splicing |
| SF3B1 | 10 | 2 | 0 | splicing factor 3b subunit 1 | Splicing |
| DAZAP1 | 2 | 4 | 2 | DAZ associated protein 1 | Splicing |
| THRAP3 | 4 | 4 | 0 | thyroid hormone receptor associated protein 3 | Splicing |
| ACVR1B | 2 | 2 | 6 | activin A receptor type 1B | TGFB signaling |
| SMAD2 | 4 | 0 | 2 | SMAD family member 2 | TGFB signaling |
| SMAD4 | 4 | 2 | 0 | SMAD family member 4 | TGFB signaling |
| ACVR1 | 4 | 0 | 0 | activin A receptor type 1 | TGFB signaling |
| DACH1 | 2 | 2 | 0 | dachshund family transcription factor 1 | TGFB signaling |

| | | | | | |
|---|---|---|---|---|---|
| ACVR2A | 2 | 0 | 0 | activin A receptor type 2A | TGFB signaling |
| TGFBR2 | 2 | 0 | 0 | transforming growth factor beta receptor 2 | TGFB signaling |
| TSC2 | 8 | 2 | 0 | TSC complex subunit 2 | TOR signaling |
| TSC1 | 2 | 0 | 0 | TSC complex subunit 1 | TOR signaling |
| MED12 | 8 | 8 | 2 | mediator complex subunit 12 | Transcription factor |
| GATA3 | 2 | 8 | 6 | GATA binding protein 3 | Transcription factor |
| FOXA1 | 4 | 6 | 4 | forkhead box A1 | Transcription factor |
| PGR | 10 | 2 | 2 | progesterone receptor | Transcription factor |
| TGIF1 | 6 | 4 | 4 | TGFB induced factor homeobox 1 | Transcription factor |
| ZNF750 | 6 | 6 | 2 | zinc finger protein 750 | Transcription factor |
| MGA | 10 | 0 | 2 | MAX dimerization protein MGA | Transcription factor |
| RUNX1 | 6 | 4 | 2 | RUNX family transcription factor 1 | Transcription factor |
| WT1 | 4 | 6 | 2 | WT1 transcription factor | Transcription factor |
| KLF5 | 6 | 2 | 2 | Kruppel like factor 5 | Transcription factor |
| MYCN | 4 | 2 | 4 | MYCN proto-oncogene, bHLH transcription factor | Transcription factor |
| EPAS1 | 4 | 2 | 2 | endothelial PAS domain protein 1 | Transcription factor |
| TBX3 | 2 | 2 | 4 | T-box transcription factor 3 | Transcription factor |
| SOX17 | 2 | 2 | 2 | SRY-box transcription factor 17 | Transcription factor |
| UNCX | 4 | 0 | 2 | UNC homeobox | Transcription factor |
| CREB3L3 | 0 | 4 | 0 | cAMP responsive element binding protein 3 like 3 | Transcription factor |
| SOX9 | 2 | 2 | 0 | SRY-box transcription factor 9 | Transcription factor |
| TAF1 | 2 | 0 | 2 | TATA-box binding protein associated factor 1 | Transcription factor |

| | | | | | |
|---|---|---|---|---|---|
| ZCCHC12 | 2 | 2 | 0 | zinc finger CCHC-type containing 12 | Transcription factor |
| CBFB | 0 | 2 | 0 | core-binding factor subunit beta | Transcription factor |
| CEBPA | 2 | 0 | 0 | CCAAT enhancer binding protein alpha | Transcription factor |
| FOXQ1 | 2 | 0 | 0 | forkhead box Q1 | Transcription factor |
| GTF2I | 2 | 0 | 0 | general transcription factor IIi | Transcription factor |
| MYC* | 0 | 2 | 0 | MYC proto-oncogene, bHLH transcription factor | Transcription factor |
| NFE2L2 | 0 | 0 | 2 | nuclear factor, erythroid 2 like 2 | Transcription factor |
| PSIP1 | 2 | 0 | 0 | PC4 and SFRS1 interacting protein 1 | Transcription factor |
| TCF12 | 0 | 2 | 0 | transcription factor 12 | Transcription factor |
| ZMYM2 | 0 | 0 | 2 | zinc finger MYM-type containing 2 | Transcription factor |
| AXIN1 | 4 | 8 | 2 | axin 1 | Wnt/B-catenin signaling |
| CTNNB1 | 6 | 2 | 4 | catenin beta 1 | Wnt/B-catenin signaling |
| TCF7L2 | 2 | 6 | 4 | transcription factor 7 like 2 | Wnt/B-catenin signaling |
| APC | 2 | 4 | 2 | APC regulator of WNT signaling pathway | Wnt/B-catenin signaling |
| AXIN2 | 0 | 0 | 2 | axin 2 | Wnt/B-catenin signaling |

* Genes previously reported mutated in CTCL
